# Comprehensive codon usage analyses of the mutations in the Parkin protein leading to Parkinsonism

**DOI:** 10.1101/2023.02.16.528897

**Authors:** Sima Biswas, Ananya Ali, Matthew Mort, Angshuman Bagchi

## Abstract

Neurodegenerative diseases typically occur due to the abnormal behavior of the proteins. One of the most prevalent neurodegenerative diseases is the Parkinson’s disease. Parkin protein encoded by PARK2 gene is the central player in the central nervous system. Mutations in Parkin lead to the onset of Parkinsonism. In this work, we made an attempt to decipher the codon usage patterns of the amino acid residues in the different domains of Parkin protein. We observed that mutations at the first two codon positions have statistically significant disease associations. This is the first such report that analyzes the correlation between the disease severity of Parkinsonism and its association with the pattern of codon usages.

## 1. Introduction

Genes are the basic functional units of heredity. They transfer the biological information from one generation to the other [1]. Genes are also responsible for the variations between organisms [2]. DNA is the basic genetic material. DNA carries the hereditary information which is passed from the parents to the offspring through the process of central dogma. DNA consists of nucleotides which determine the genetic codes necessary to generate the amino acids. The genetic codes are redundant and the same genetic code may be used for generating a number of different amino acids. Only two amino acids, methionine and tryptophan, are encoded by a single type of codon [3]. It is, however, observed that there are specificities of codon usage patterns for a particular amino acid and it varies between organisms [4-7]. Hence, changes during DNA transcription might change the genetic code which eventually alters the mode of protein synthesis. Single nucleotide substitutions (SNS) in the codons induces genetic mutations. Base substitutions, base alterations, insertions, deletions can have a variety of effects [8]. Only in case of silent mutations the base substitutions cannot affect the protein product directly but could impact the protein sequence by disruption to pre-mRNA splicing. On the other hand, single nucleotide substitutions which alter the codon may generate missense mutations (amino acid substitution) which could result in altered protein functions thereby leading to deleterious effects [9]. As in some cases multiple codons can encode the same amino acid, it was observed that changes in the first two nucleotide positions of a codon are mostly likely to alter the encoded amino acid, with the 3^rd^ nucleotide position being referred to as the wobble position. Mutations within codons, may alter protein primary sequence and consequently may alter gene functions which subsequently may lead to disease or disease susceptibility. In this work, we focus on the codon usage patterns in the Parkin protein, which is the central player in different biological pathways. Mutations in the Parkin protein often lead to the onset of Parkinsonism [10] Many mutations reported so far in the various Database. Therefore, it may be speculated that any change in the codon usage pattern of the PRKN gene, which codes of the Parkin protein, might have some effect on the functionality of the protein thereby leading to the onset of Parkinsonism.

One of the very common and severe neurodegenerative disorders is the Parkinson’s disease (PD). The loss of substantial dopaminergic neurons is the main cause of the onset of PD [11]. Environmental and genetic factors are responsible for the pathogenesis of this multifactorial disease [12]. Sporadic and familial are the two groups of the disease form. PARK2, PINK1 and DJ1 gene mutations are responsible for the early onset of Parkinson disease [13]. Parkin is a multi-domain protein containing the domains referred to as Ubl, L-region, RING0, RING1, Rep, IBR and RING2 [14]. More than 150 mutations are spread throughout the different domains of the Parkin protein and these mutations lead to the onset of Parkinson’s disease [15]. In this current work, we first collected the mutations in the different domains of Parkin protein from different databases. We used in-silico tools to categorize the mutations as per their severities. We then used different statistical tests to correlate the patterns of the changes in the codon positions and their relationships with the nature of the mutations. With this work, we tried to predict how a particular change in a specific codon position could bring about the changes in the Parkin protein and how that change could influence the disease onset.

## 2. Materials & Methods

### 2.1 Collection of mutations

The information pertaining to the mutations in the human Parkin protein were collected from literature, PDmutDB, Genome Aggregation Database (https://gnomad.broadinstitute.org/) and COSMIC databases. The Parkinson’s disease Mutation Database or PDmutDB (https://www.molgen.vib-ua.be/PDMutDB/) is a store-house of all the mutation related data associated with PD. We also searched in COSMIC database (http://cancer.sanger.ac.uk/) which is mainly related to somatic gene mutations leading to different types of cancers. However, there are certain mutations which appear in both cancer and Parkinson’s disease. Therefore, we used this database in order to get a comprehensive mutation data.

### 2.2 Prediction of pathogenicity of mutations

The next target was to assign a severity score to the collected mutations. For that, we used the same principle as we used in our previous work [16]. We used the following servers:

a. SNAP2 (Scalable Nucleotide Alignment Program) (https://www.rostlab.org/services/snap/): The tool is a machine learning based server. It utilizes the principles of “Neural network”. The tool takes the amino acid sequence as input along-with the information of the mutations. The tool classifies the mutations on the basis of evolutionary and other relevant information as deleterious or neutral [17].
b. PolyPhen 2.0 (Polymorphism phenotyping 2.0, http://genetics.bwh.harvard.edu/pph2/): PolyPhen 2.0 uses physical and comparative approaches for the prediction of the effect of the amino acid substitution and their impact on protein structure. An input of the polyphen2 is the UniProt accession number/FASTA sequence and the details of amino acids substitutions. Based on inputs and evolutionary conservation of amino acid the predictions are being made [18].
c. SDM (Site directed mutator, http://marid.bioc.cam.ac.uk/sdm2/prediction): SDM is a statistical approach for analyzing amino acid substitutions. Known 3D structures are to be given as inputs to the server. The mutational effects are determined by the study of the details of the amino acid residues in homologous proteins [19].
d. PROVEAN (Protein variation effect analyzer, http://provean.jcvi.org/index.php): PROVEAN, a homology-based prediction tool, analyses the impact of an amino acid substitution in the protein structure. Based on the scores, the mutations are labeled as deleterious or neutral [20].
e. SIFT (Sorting Intolerant from Tolerant amino acid substitutions, https://sift.bii.a-star.edu.sg/): Score from multiple sequence alignment is used by SIFT for the predictions of the effects of amino acid substitutions. SIFT scores span the range starting from 0 to 1. The damaging amino acid substitutions are those which get scores below 0.05 [21].
f. Align GV (GranthamVariation) GD (GranthamDeviation) (http://agvgd.iarc.fr/agvgd): The tool Align GV-GD analyzes physico-chemical properties and changes in biophysical characteristics due to amino acid substitutions [22].
g. SNPs&GO (https://snps-andgo.biocomp.unibo.it/snps-and-go/): The support vector machine (SVM) based prediction method SNPs&GO predicts disease associated or neutral mutations from the amino acid sequences of proteins [23].

Then we analyzed the properties of the mutations as obtained from the aforementioned servers. We scored the mutations on a scale of 0-7 based on the prediction results, where 7 being the most damaging mutation and 0 being the tolerated mutation. The classification scheme employed in this work is presented in table1:

**Table1:** Classifications of the mutations’ pathogenicity

| Score | Classification of mutation | Details |
| --- | --- | --- |
| 7 | Damaging or Pathogenic | All the 7 servers used predicted the mutation to be damaging |
| 6 | Damaging or Pathogenic | Out of the 7 servers used, 6 of the servers predicted the mutation to be damaging |
| 3 to 5 | Moderate | Out of the 7 servers used, 3 to 5 servers predicted the mutation to be damaging and the rest of the servers predicted the mutation to be tolerable. |
| 0 to 2 | Least or non-damaging / non-disease causing | Out of the 7 servers used, most of the servers predicted the mutation to be tolerable. |

### 2.3 Compositional Properties

The compositional properties were analyzed by measuring the percentages of the bases present in the genome. The percentages of A, T, G and C were calculated before and after the mutation. Position wise the percentages of the aforementioned nucleotide bases were calculated. We also counted the percentages of the base changes observed after mutations.

### 2.4 Statistical Analyses

We arranged our raw data under three broad variables, viz., the domain specificity, the codon position of mutations and severity of the disease. The variable domain specificity had 7 sub-variables based on the distributions of the amino acids in the Parkin protein. The sub-variables were the domains, viz., Ubl, L-region, RING0, RING1, Rep, IBR and RING2 in the Parkin protein. The second variable, codon position, was expressed by a discrete numerical value pointing towards the presence of the mutation on the positions of the triplet codon. The value of the second variable was identified by looking into the dispositions of the nucleotides in the wild type and mutant codons. The third variable was referred to as the disease severity due to the mutation. This variable was determined using the aforementioned 7 different servers. In order to determine the relationship between these factors, we performed a homogeneity chi squared test using the PAST software [24-25]. We calculated the correlations between

a. the position of the mutation with its effect on disease onset
b. identity of the domain where the mutation is present with its effect on disease onset

The results were presented as mutation position in the codon with the severity of the disease and domain identity for each position with the severity of the disease.

Furthermore, we performed the aforementioned analyses on a set of neutral gene mutations as obtained from Genome Aggregation Database. This was done to make a comparison between the different types of disease associated mutations with the neutral variants. The set of neutral variants is used as a control dataset.

## 3. Results and Discussions

### 3.1 Prediction of pathogenicity

We collected total of 252 missense and 144 synonymous substitutions in the different domains of the Parkin protein. We then analyzed the mutations using 7 different servers to check the effects of the mutations on disease onset. From the aforementioned analysis, following domain specific results were obtained and they were presented in table2.

**Table2:** Domain wise distributions of the mutations and their patterns

| Name of the domain | Type of mutations |  |  |  |
| --- | --- | --- | --- | --- |
|  | Damaging | Moderate | Tolerated | Synonymous<br>(Control) |
| Ubl | 16 | 11 | 13 | 17 |
| L- Region | 2 | 8 | 22 | 17 |
| RING0 | 19 | 22 | 2 | 21 |
| RING1 | 28 | 26 | 11 | 41 |
| REP | 4 | 10 | 6 | 8 |
| IBR | 11 | 13 | 4 | 21 |
| RING2 | 16 | 8 | 2 | 19 |

Ubl domain: In this domain there were 40 mutations present. Out of the 40 mutations, 16 mutations were predicted to be damaging.

L-regions. Out of the total 32 mutations present in this domain, 2 mutations were found to be damaging.

RING0: Out of the total 43 mutations present in this domain, 19 mutations were found to be damaging.

RING1: Out of the total 65 mutations present in this domain, 28 mutations were found to be damaging.

REP domain: In this domain there were 20 mutations present. Out of the 20 mutations, 4 mutations were predicted to be damaging.

IBR domain: 28 mutations were present in this domain. Out the total number of mutations present in the domain, 11 mutations were predicted to be damaging.

RING2: The domain contains a total of 26 mutations. Among them 16 mutations were predicted to be damaging.

We computed the number of synonymous substitutions at the corresponding domains of Parkin protein (table2). These synonymous substitutions were used as control dataset.

It was observed from the analysis that the RING1 domain has the most damaging mutations. RING1 domain is known to bind to the E2 Ubiquitin conjugating enzyme [14]. Each and every domain of the Parkin protein is important. The Ubl domain in Parkin protein interacts with PINK1 protein and this interaction is needed for the activation of the Parkin protein. Catalytic cystine residues are located in the RING2 domain. Other domains in the Parkin protein interact with each other to maintain the structural integrity of the protein. The mutations in these domains lead to the impairment of the functionality of the domain and thereby the overall activity of the Parkin protein.

### 3.2 Effect of base compositions in the codons

The nucleotide base composition offers a diverse choice for codon selection. Therefore, we computed the percentages of the different bases at the respective positions of the codon in the different domains of the Parkin protein. We analyzed the base compositions of the different domains of the Parkin protein for the different neutral variants which were being used as controls. The results were given in the tables 3.1 to 3.7.

**Table 3.1 A:**
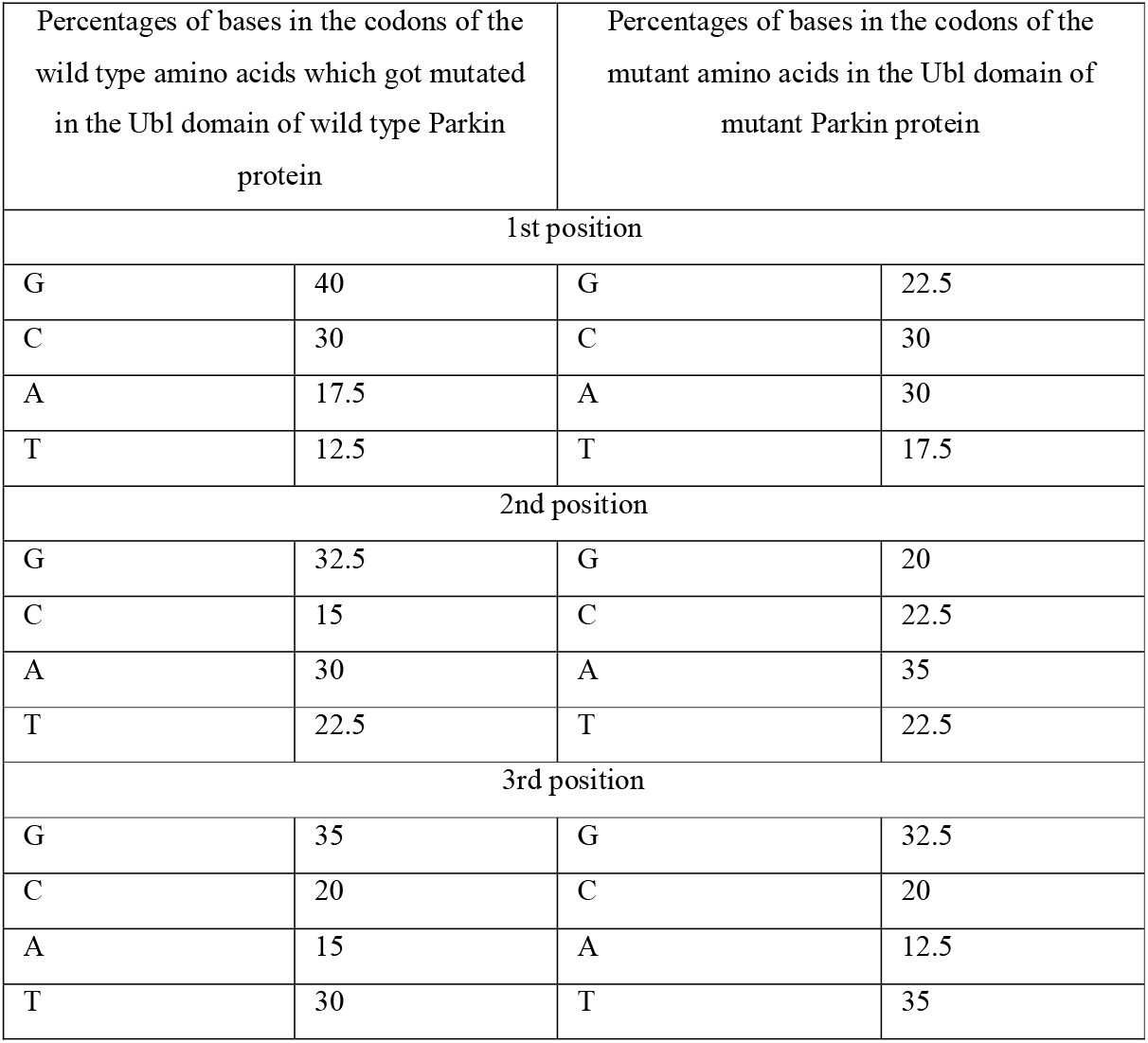
Analyses of the base compositions of the codons of the wild type and mutant amino acids in the Ubl domain of Parkin protein

| Percentages of bases in the codons of the wild type amino acids which got mutated in the Ubl domain of wild type Parkin protein |  | Percentages of bases in the codons of the mutant amino acids in the Ubl domain of mutant Parkin protein |  |
| --- | --- | --- | --- |
| 1st position |  |  |  |
| G | 40 | G | 22.5 |
| C | 30 | C | 30 |
| A | 17.5 | A | 30 |
| T | 12.5 | T | 17.5 |
| 2nd position |  |  |  |
| G | 32.5 | G | 20 |
| C | 15 | C | 22.5 |
| A | 30 | A | 35 |
| T | 22.5 | T | 22.5 |
| 3rd position |  |  |  |
| G | 35 | G | 32.5 |
| C | 20 | C | 20 |
| A | 15 | A | 12.5 |
| T | 30 | T | 35 |

**Table 3.1 B:** Analyses of the base compositions of the codons of the wild type and mutant amino acids in the Ubl domain of Parkin protein in cases of neutral variants

| Percentages of bases in the codons of the wild type amino acids which got mutated in the Ubl domain of wild type Parkin protein |  | Percentages of bases in the codons of the mutant amino acids in the Ubl domain of mutant Parkin protein |  |
| --- | --- | --- | --- |
| 1st position |  |  |  |
| G | 29.41 | G | 29.41 |
| C | 41.17 | C | 41.17 |
| A | 23.53 | A | 23.53 |
| T | 5.88 | T | 5.88 |
| 2nd position |  |  |  |
| G | 23.53 | G | 23.53 |
| C | 17.65 | C | 17.65 |
| A | 35.29 | A | 35.29 |
| T | 23.53 | T | 23.53 |
| 3rd position |  |  |  |
| G | 58.82 | G | 11.76 |
| C | 23.53 | C | 11.76 |
| A | 11.76 | A | 52.94 |
| T | 5.88 | T | 23.53 |

**Table 3.2 A:** Analyses of the base compositions of the codons of the wild type and mutant amino acids in the L-Region of Parkin protein

| Percentages of bases in the codons of the wild type amino acids which got mutated in the L-Region of wild type Parkin protein |  | Percentages of bases in the codons of the mutant amino acids in the L-Region of mutant Parkin protein |  |
| --- | --- | --- | --- |
| 1st position |  |  |  |
| G | 40.62 | G | 18.75 |
| C | 37.5 | C | 31.25 |
| A | 15.62 | A | 37.5 |
| T | 6.25 | T | 12.5 |
| 2nd position |  |  |  |
| G | 25 | G | 21.87 |
| C | 37.5 | C | 18.75 |
| A | 25 | A | 43.75 |
| T | 12.5 | T | 15.62 |
| 3rd Position |  |  |  |
| G | 34.37 | G | 28.12 |
| C | 34.37 | C | 34.37 |
| A | 25 | A | 25 |
| T | 6.25 | T | 12.5 |

**Table 3.2 B:** Analyses of the base compositions of the codons of the wild type and mutant amino acids in the L-Region of Parkin protein in cases of neutral variants

| Percentages of bases in the codons of the wild type amino acids which got mutated in the L-Region of wild type Parkin protein |  | Percentages of bases in the codons of the mutant amino acids in the L-Region of mutant Parkin protein |  |
| --- | --- | --- | --- |
| 1st position |  |  |  |
| G | 52.94 | G | 52.94 |
| C | 29.41 | C | 17.64 |
| A | 11.76 | A | 11.76 |
| T | 5.88 | T | 17.64 |
| 2nd position |  |  |  |
| G | 11.76 | G | 11.76 |
| C | 29.41 | C | 29.41 |
| A | 35.29 | A | 35.29 |
| T | 23.52 | T | 23.52 |
| 3rd Position |  |  |  |
| G | 64.70 | G | 11.76 |
| C | 23.52 | C | 29.41 |
| A | 0 | A | 29.41 |
| T | 11.76 | T | 29.41 |

**Table 3.3 A:** Analyses of the base compositions of the codons of the wild type and mutant amino acids in the RING0 domain of Parkin protein

| Percentages of bases in the codons of the wild type amino acids which got mutated in the RING0 domain of wild type Parkin protein |  | Percentages of bases in the codons of the mutant amino acids in the RING0 domain of mutant Parkin protein |  |
| --- | --- | --- | --- |
| 1st position |  |  |  |
| G | 30.95 | G | 16.66 |
| C | 28.57 | C | 21.42 |
| A | 19.04 | A | 26.19 |
| T | 21.42 | T | 35.71 |
| 2nd position |  |  |  |
| G | 42.85 | G | 14.28 |
| C | 23.80 | C | 11.90 |
| A | 19.04 | A | 40.47 |
| T | 14.28 | T | 33.33 |
| 3rd position |  |  |  |
| G | 23.80 | G | 21.42 |
| C | 30.95 | C | 30.95 |
| A | 28.57 | A | 23.80 |
| T | 16.66 | T | 23.80 |

**Table 3.3 B:** Analyses of the base compositions of the codons of the wild type and mutant amino acids in the RING0 domain of Parkin protein in cases of neutral variants

| Percentages of bases in the codons of the wild type amino acids which got mutated in the RING0 domain of wild type Parkin protein |  | Percentages of bases in the codons of the mutant amino acids in the RING0 domain of mutant Parkin protein |  |
| --- | --- | --- | --- |
| 1st position |  |  |  |
| G | 28.57 | G | 28.57 |
| C | 33.33 | C | 33.33 |
| A | 23.80 | A | 23.80 |
| T | 14.28 | T | 14.28 |
| 2nd position |  |  |  |
| G | 23.80 | G | 23.80 |
| C | 38.09 | C | 38.09 |
| A | 19.04 | A | 19.04 |
| T | 19.04 | T | 19.04 |
| 3rd position |  |  |  |
| G | 19.04 | G | 14.28 |
| C | 52.38 | C | 19.04 |
| A | 9.52 | A | 33.33 |
| T | 19.04 | T | 33.33 |

**Table 3.4 A:** Analyses of the base compositions of the codons of the wild type and mutant amino acids in the RING1 domain of Parkin protein

| Percentages of bases in the codons of the wild type amino acids which got mutated in the RING1 domain of wild type Parkin protein |  | Percentages of bases in the codons of the mutant amino acids in the RING1 domain of mutant Parkin protein |  |
| --- | --- | --- | --- |
| 1st position |  |  |  |
| G | 33.33 | G | 18.18 |
| C | 31.81 | C | 24.24 |
| A | 21.21 | A | 30.30 |
| T | 13.63 | T | 27.27 |
| 2nd position |  |  |  |
| G | 30.30 | G | 27.27 |
| C | 19.69 | C | 13.63 |
| A | 22.72 | A | 28.78 |
| T | 27.27 | T | 30.30 |
| 3rd position |  |  |  |
| G | 31.81 | G | 27.27 |
| C | 31.81 | C | 33.33 |
| A | 15.15 | A | 13.63 |
| T | 21.21 | T | 25.75 |

**Table 3.4 B:**
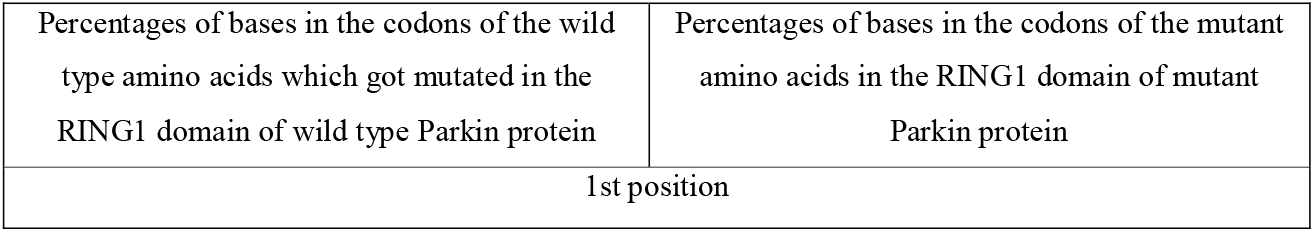
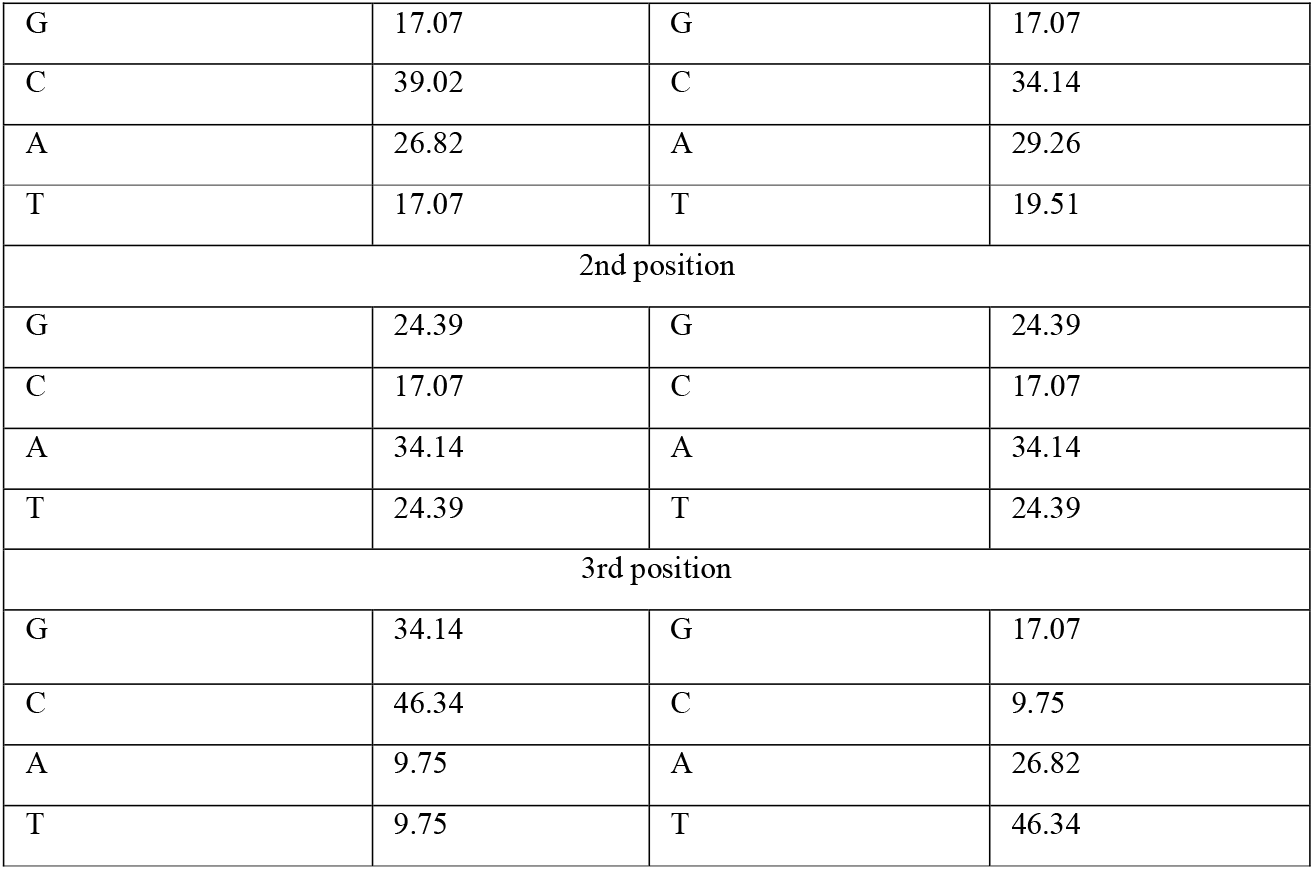
Analyses of the base compositions of the codons of the wild type and mutant amino acids in the RING1 domain of Parkin protein in cases of neutral variants

| Percentages of bases in the codons of the wild type amino acids which got mutated in the RING1 domain of wild type Parkin protein | Percentages of bases in the codons of the mutant amino acids in the RING1 domain of mutant Parkin protein |
| --- | --- |
| 1st position |  |

|  |  |  |  |
| --- | --- | --- | --- |
| G | 17.07 | G | 17.07 |
| C | 39.02 | C | 34.14 |
| A | 26.82 | A | 29.26 |
| T | 17.07 | T | 19.51 |
| 2nd position |  |  |  |
| G | 24.39 | G | 24.39 |
| C | 17.07 | C | 17.07 |
| A | 34.14 | A | 34.14 |
| T | 24.39 | T | 24.39 |
| 3rd position |  |  |  |
| G | 34.14 | G | 17.07 |
| C | 46.34 | C | 9.75 |
| A | 9.75 | A | 26.82 |
| T | 9.75 | T | 46.34 |

**Table 3.5 A:** Analyses of the base compositions of the codons of the wild type and mutant amino acids in the REP domain of Parkin protein

| Percentages of bases in the codons of the wild type amino acids which got mutated in the REP domain of wild type Parkin protein |  | Percentages of bases in the codons of the mutant amino acids in the REP domain of mutant Parkin protein |  |
| --- | --- | --- | --- |
| 1st position |  |  |  |
| G | 55 | G | 30 |
| C | 10 | C | 15 |
| A | 20 | A | 40 |
| T | 15 | T | 15 |
| 2nd position |  |  |  |
| G | 20 | G | 30 |
| C | 35 | C | 20 |
| A | 35 | A | 30 |
| T | 10 | T | 20 |
| 3rd position |  |  |  |
| G | 5 | G | 0 |
| C | 35 | C | 40 |
| A | 40 | A | 35 |
| T | 20 | T | 25 |

**Table 3.5 B:** Analyses of the base compositions of the codons of the wild type and mutant amino acids in the REP domain of Parkin protein in cases of neutral variants

| Percentages of bases in the codons of the wild type amino acids which got mutated in the REP domain of wild type Parkin protein |  | Percentages of bases in the codons of the mutant amino acids in the REP domain of mutant Parkin protein |  |
| --- | --- | --- | --- |
| 1st position |  |  |  |
| G | 87.5 | G | 87.5 |
| C | 0 | C | 0 |
| A | 12.5 | A | 12.5 |
| T | 0 | T | 0 |
| 2nd position |  |  |  |
| G | 0 | G | 0 |
| C | 62.5 | C | 62.5 |
| A | 12.5 | A | 12.5 |
| T | 25 | T | 25 |
| 3rd position |  |  |  |
| G | 0 | G | 37.5 |
| C | 75 | C | 12.5 |
| A | 12.5 | A | 0 |
| T | 12.5 | T | 50 |

**Table 3.6 A:** Analyses of the base compositions of the codons of the wild type and mutant amino acids in the IBR domain of Parkin protein

| Percentages of bases in the codons of the wild type amino acids which got mutated in the IBR domain of wild type Parkin protein |  | Percentages of bases in the codons of the mutant amino acids in the IBR domain of mutant Parkin protein |  |
| --- | --- | --- | --- |
| 1st position |  |  |  |
| G | 42.85 | G | 21.42 |
| C | 35.71 | C | 32.14 |
| A | 10.71 | A | 25 |
| T | 10.71 | T | 21.42 |
| 2nd Position |  |  |  |
| G | 39.28 | G | 21.42 |
| C | 25 | C | 14.28 |
| A | 25 | A | 39.28 |
| T | 10.71 | T | 25 |
| 3rd Position |  |  |  |
| G | 35.71 | G | 28.57 |
| C | 32.14 | C | 32.14 |
| A | 10.71 | A | 21.42 |
| T | 21.42 | T | 17.85 |

**Table 3.6 B:** Analyses of the base compositions of the codons of the wild type and mutant amino acids in the IBR domain of Parkin protein in cases of neutral variants

| Percentages of bases in the codons of the wild type amino acids which got mutated in the IBR domain of wild type Parkin protein |  | Percentages of bases in the codons of the mutant amino acids in the IBR domain of mutant Parkin protein |  |
| --- | --- | --- | --- |
| 1st position |  |  |  |
| G | 52.38 | G | 52.38 |
| C | 28.57 | C | 28.57 |
| A | 4.76 | A | 4.76 |
| T | 14.28 | T | 14.28 |
| 2nd Position |  |  |  |
| G | 47.61 | G | 47.61 |
| C | 19.04 | C | 19.04 |
| A | 14.28 | A | 14.28 |
| T | 19.04 | T | 19.04 |
| 3rd Position |  |  |  |
| G | 47.61 | G | 14.28 |
| C | 47.61 | C | 4.76 |
| A | 4.76 | A | 47.61 |
| T | 0 | T | 33.33 |

**Table 3.7 A:** Analyses of the base compositions of the codons of the wild type and mutant amino acids in the RING2 domain of Parkin protein

| Percentages of bases in the codons of the wild type amino acids which got mutated in the RING2 domain of wild type Parkin protein |  | Percentages of bases in the codons of the mutant amino acids in the RING2 domain of mutant Parkin protein |  |
| --- | --- | --- | --- |
| 1st position |  |  |  |
| G | 30.76 | G | 15.38 |
| C | 30.76 | C | 38.46 |
| A | 11.53 | A | 26.92 |
| T | 26.92 | T | 19.23 |
| 2nd position |  |  |  |
| G | 46.15 | G | 23.07 |
| C | 15.38 | C | 3.84 |
| A | 26.92 | A | 34.61 |
| T | 11.53 | T | 38.46 |
| 3rd position |  |  |  |
| G | 30.76 | G | 30.76 |
| C | 50 | C | 50 |
| A | 7.69 | A | 7.69 |
| T | 11.53 | T | 11.53 |

**Table 3.7 B:** Analyses of the base compositions of the codons of the wild type and mutant amino acids in the RING2 domain of Parkin protein in cases of neutral variants

| Percentages of bases in the codons of the wild type amino acids which got mutated in the RING2 domain of wild type Parkin protein |  | Percentages of bases in the codons of the mutant amino acids in the RING2 domain of mutant Parkin protein |  |
| --- | --- | --- | --- |
| 1st position |  |  |  |
| G | 42.10 | G | 42.10 |
| C | 42.10 | C | 42.10 |
| A | 10.52 | A | 10.52 |
| T | 5.26 | T | 5.26 |
| 2nd position |  |  |  |
| G | 15.78 | G | 15.78 |
| C | 26.31 | C | 26.31 |
| A | 15.78 | A | 15.78 |
| T | 42.10 | T | 42.10 |
| 3rd position |  |  |  |
| G | 15.78 | G | 15.78 |
| C | 63.15 | C | 15.78 |
| A | 15.78 | A | 10.52 |
| T | 5.26 | T | 57.89 |

It was observed that the percentages of the bases were different in the different regions. It was previously reported that there could be a strong correlation between the GC content at the third codon position with the usages of codons. This pattern of codon usage is associated with the amounts of proteins and mRNAs in cells [26]. From the aforementioned tables it was apparent that the GC contents at the third codon positions for the amino acids belonging to the different domains would vary significantly in both the wild type and mutant Parkin proteins. The aforementioned GC content was found to be the maximum for the amino acids of the RING2 domain. The RING2 domain of Parkin has a very significant role. It was also reported that the single RING2 domain of Parkin would be able to exert the catalytic activity of the entire Parkin protein even in absence of the other domains [27]. Therefore, the high GC content at the third codon position for the amino acids in the RING2 domain would point towards the probable ease of synthesis of the domain due to its biological activities.

We then tried to analyze the domain-wise changes in the base compositions in the codons of the amino acids in case of neutral variants. We performed this analysis as a control experiment. Only for the RING1 domain there were changes in base compositions at both the first and third codon positions whereas for all other domains changes in base compositions were observed only at the third codon position. Surprisingly, the domain wise base compositions in all the positions were found to vary when we tried to compare the same for the non-neutral mutations. In other words, the distributions of nucleotides were found to be different in the codon positions of the wild type and mutant amino acids for those non-neutral mutations irrespective of the severity of the disease onset. This is contrary to the fact that for neutral mutations, base compositions at only the third codon positions would vary except for RING1 domain.

### 3.3 Correlation between the position of the mutation with its effect on disease onset

We used statistical correlation analysis on the collected data to check if there is any influence of the position of the mutation on the codon with the severity of the mutations. We performed the analysis on the entire dataset containing the mutations from all the domains of the Parkin protein. It was observed that there was a no significant association between the position of the mutation on the codon and severity of the disease with a p-value of 0.26638. However, our analyses revealed that the onset of Parkinsonism is more prevalent when mutations would appear at codon position 1 or 2. This trend was observed for all the aforementioned domains of the Parkin protein. On the other hand, mutations at codon position 3 do not have any significant relationship with the severity of the disease. On further analysis, it was observed that the effects of mutations at the codon position 1 [p (association): 0.0099023] and 2 [p (association): 0.000239] have more heterogeneous outcomes than at codon position 3 [p (association): 0.3273]. Statistical analyses of our data showed that mutations at codon positions 1 and 2 are strongly associated with the severity of the Parkinson’s disease as compared to codon position 3.

We then computed the relationships between the changes in base compositions at the respective codon positions in disease associated and neutral mutations. The comparison between neutral mutations and disease associated mutations would show that a significant difference (with a p-value < 0.000001) would be present among the two types of mutations in their codon preferences. While Neutral mutations were found to be mostly located at third codon positions, disease associated mutations showed more preferences towards positions 1 and 2.

## Conclusion

This is the first work on the association of the appearance of mutations on the codon positions with the severity of disease onset. The basic aim of the work was to come up with the prediction of plausible mechanism of disease onset. From our analyses, we could safely conclude that if the first two positions of the codon get mutated the extent of Parkinsonism would be more as compared to the third position. However, for neutral variants, changes were observed only at the third codon position whereas the first two codon positions remained mostly unaffected.

## Acknowledgement

The authors acknowledge the financial support from the DBT funded Bioinformatics Infrastructure Facility Center (Sanction no.: BT/PR40162/BTIS/137/48/2022 sanctioned to Prof. Angshuman Bagchi) of University of Kalyani for support. Sima Biswas receives funding from University of Kalyani as the University Research Scholar.

## Author Contributions

AB and MM conceptualized the work. SB and AA performed the analysis. All the authors were involved in writing the manuscript. All the authors agreed to submit the manuscript.

