## Supplementary for "Comprehensive codon usage analyses of the mutations in the Parkin protein leading to Parkinsonism"

Table 1.1Mutation analysis results of Ubl region

| CODON | POSITION | CODON CHANGE | NEW CODON | **MUTATION** | **GV/GD** | **SIFT** | **PolyPhen2** | **SNPs&GO** | **SNAP** | **PROVEN** | **SDM** | **SCORE** | **INFERENCE** |
| --- | --- | --- | --- | --- | --- | --- | --- | --- | --- | --- | --- | --- | --- |
| ATA | 2 | A>G | GAT | I2V | Class C25 | Tolerated | Benign | Neutral | Neutral | Neutral | Increased | 0 out of 7 | LOW |
| AGG | 6 | G>T | AGT | R6S | Class C65 | AFFECT PROTEIN FUNCTION | PROBABLY DAMAGING | Disease | effect | Deleterious | Reduced stability | 7 out of 7 | HIGH |
| TCC | 9 | T>G | GCC | S9A | Class C65 | Tolerated | Probably Damaging | Neutral | Neutral | Neutral | Increased | 2 out of 7 | LOW |
| AGC | 10 | G>A | AAC | S10N | Class C45 | Tolerated | Benign | Neutral | Neutral | Neutral | Increased | 1 out of 7 | LOW |
| GGT | 12 | GG>TT | TTT | G12F | Class C65 | Affect protein function | Probably Damaging | Neutral | Effect | Deleterious | Reduced | 6 out of 7 | HIGH |
| GGT | 12 | G>C | CGT | G12R | Class C65 | Tolerated | Probably Damaging | Neutral | Effect | Neutral | Reduced | 4 out of 7 | MODERATE |
| TTC | 13 | T>G | GTC | F13V | Class C45 | Tolerated | Possibly Damaging | Neutral | Neutral | Neutral | Reduced | 3 out of 7 | modarate |
| CCA | 14 | C>T | TCA | P14S | Class C65 | Tolerated | Probably Damaging | Neutral | Neutral | Deleterious | Increased | 3 out of 7 | LOW |
| GTG | 15 | G>A | ATG | V15M | Class C15 | Affect protein function | Probably Damaging | Neutral | Effect | Neutral | Reduced | 4 out of 7 | HIGH |
| GTC | 17 | T>C | GCC | F13V | Class C45 | Tolerated | Possibly Damaging | Neutral | Neutral | Neutral | Reduced | 3 out of 7 | 1^st^ |
| GAT | 18 | G>C | CAT | D18H | Class C65 | Tolerated | Benign | Neutral | Neutral | Deleterious | Increased | 2 out of 7 | LOW |
| GAT | 18 | G>A | AAT | D18N | Class C15 | Affect protein function | Benign | Neutral | Neutral | Neutral | Reduced | 2 out of 7 | LOW |
| AGC | 22 | G>T | ATC | S22I | Class C65 | Affect protein function | Probably Damaging | Neutral | Effect | Deleterious | Increased | 5 out of 7 | MODERATE |
| ATC | 23 | A>G | GTC | I23V | Class C25 | Tolerated | Benign | Neutral | Neutral | Neutral | Reduced | 1 out of 7 | LOW |
| TTC | 24 | T>C | TCC | F24S | Class C65 | Tolerated | Benign | Neutral | Neutral | Neutral | Reduced | 2 out of 7 | LOW |
| GAG | 28 | G>A | AAG | E28K | Class C55 | Affect protein function | Probably Damaging | Disease | Effect | Deleterious | Reduced | 7 out of 7 | High |
| GTG | 29 | T>C | GCG | V29A | Class C55 | Tolerated | Benign | Neutral | Neutral | Neutral | Increased | 1 out of 7 | LOW |
| GCT | 31 | C>A | GAT | A31D | Class C65 | Affect protein function | Probably Damaging | Disease | Effect | Deleterious | Reduced | 7 out of 7 | HIGH |
| CGA | 33 | G>A | CAA | R33Q | Class C35 | Tolerated | Possibly Damaging | Neutral | Neutral | Neutral | Increased | 1 out of 7 | LOW |
| CAG | 34 | A>G | CGG | Q34R | Class C35 | Affect protein function | Probably Damaging | Neutral | Effect | Neutral | Reduced | 4 out of 7 | Modarate |
| CCG | 37 | C>T | CTG | P37L | Class C65 | Affect protein function | Probably Damaging | Neutral | Effect | Deleterious | Reduced | 6 out of 7 | HIGH |
| CGT | 42 | C>T | TGT | R42S | Class C65 | Affect protein function | Probably Damaging | Disease | Effect | Deleterious | Reduced | 7 out of 7 | HIGH |
| CGT | 42 | G>A | CAT | R42C | Class C65 | Tolerated | Possibly Damaging | Disease | Neutral | Deleterious | Reduced | 5 out of 7 | MODERATE |
| CGT | 42 | G>C | CCT | R42H | Class C25 | Tolerated | Benign | Neutral | Effect | Deleterious | Reduced | 3 out of 7 | Modarate |
| CGT | 42 | C>A | AGT | R42P | Class C65 | Affect protein function | Probably Damaging | Disease | Effect | Deleterious | Reduced | 7 out of 7 | HIGH |
| GTG | 43 | G>T | TTG | V43L | Class C25 | Affect protein function | Possibly Damaging | Neutral | Effect | Neutral | Reduced | 4 out of 7 | Modarate |
| GCA | 46 | G>C | CCA | A46P | Class C25 | Affect protein function | Probably Damaging | Disease | Effect | Deleterious | Reduced | 6 out of 7 | HIGH |
| GCA | 46 | G>A | ACA | A46T | Class C55 | Affect protein function | Probably Damaging | Disease | Effect | Deleterious | Reduced | 7 out of 7 | HIGH |
| GGG | 47 | G>C | CGG | G47R | Class C65 | Affect protein function | Probably Damaging | Disease | Effect | Deleterious | Reduced | 7 out of 7 | HIGH |
| GAG | 49 | G>A | AAG | E49K | Class C55 | Affect protein function | Probably Damaging | Disease | Effect | Deleterious | Reduced | 7 out of 7 | HIGH |
| TGG | 54 | T>A | AGG | W54R | Class C65 | Tolerated | Benign | Neutral | Effect | Neutral | Reduced | 3 out of 7 | Modaarte |
| GTG | 56 | T>A | GAG | V56E | Class C65 | Affect protein function | Possibly Damaging | Neutral | Effect | Deleterious | Reduced | 6 out of 7 | HIGH |
| AAT | 58 | A>T | ATT | N58I | Class C65 | Tolerated | Benign | Neutral | Effect | Deleterious | Increased | 3 out of 7 | Modarate |
| TGT | 59 | G>T | TTT | C59F | Class C65 | Affect protein function | Probably Damaging | Disease | Effect | Deleterious | Reduced | 7 out of 7 | HIGH |
| GAT | 62 | G>C | CAT | D62H | Class C65 | Tolerated | Possibly Damaging | Neutral | Neutral | Neutral | Increased | 2 out of 7 | LOW |
| CAG | 63 | C>A | AAG | Q63K | Class C45 | Affect protein function | Possibly Damaging | Disease | Effect | Neutral | Reduced | 6 out of 7 | HIGH |
| CAC | 68 | C>T | TAC | H68Y | Class C65 | Affect protein function | Probably Damaging | Disease | Effect | Deleterious | Increased | 6 out of 7 | HIGH |
| CAG | 71 | A>C | CCG | Q71P | Class C65 | Tolerated | Benign | Neutral | Effect | Neutral | Reduced | 3 out of 7 | Modarate |
| CAG | 71 | A>G | CGG | Q71R | Class C35 | Tolerated | Benign | Neutral | Neutral | Neutral | Increased | 0 out of 7 | LOW |
| AAA | 76 | A>G | GAA | K76E | Class C55 | Tolerated | Benign | Neutral | Neutral | Neutral | Reduced | 2 out of 7 | LOW |

Table 1.2: Mutation analysis results of L region

| CODON | position | CODON CHANGE | NEW CODON | MUTANT | proven | GV/GD | SIFT | SNAP | SDM | POLYPHEN2 | SNPs & GO | SCORE | INFERENCE |
| --- | --- | --- | --- | --- | --- | --- | --- | --- | --- | --- | --- | --- | --- |
| GCA | 82 | C>A | GAA | A82E | Neutral | Class C65 | AFFECT PROTEIN FUNCTION | effect | Reduced stability | BENIGN | Neutral | 4 | modarate |
| ACT | 83 | A>G | GCT | T83A | Neutral | Class C55 | AFFECT PROTEIN FUNCTION | neutral | Increased stability | BENIGN | Neutral | 2 | low |
| GAC | 86 | G>A | AAC | D86N | Neutral | Class C15 | AFFECT PROTEIN FUNCTION | neutral | Increased stability | BENIGN | Neutral | 1 | low |
| GAC | 87 | G>T | TAC | D87Y | Neutral | Class C65 | AFFECT PROTEIN FUNCTION | neutral | Increased stability | BENIGN | Neutral | 2 | low |
| CCC | 88 | C>G | CGC | P88R | Neutral | Class C65 | AFFECT PROTEIN FUNCTION | neutral | Increased stability | POSSIBLY DAMAGING | Neutral | 3 | modarate |
| CCC | 88 | C>T | CTC | P88L | Neutral | Class C65 | AFFECT PROTEIN FUNCTION | neutral | Increased stability | BENIGN | Neutral | 2 | low |
| AGA | 89 | A>G | GGA | R89G | Neutral | Class C65 | AFFECT PROTEIN FUNCTION | effect | Reduced stability | BENIGN | Neutral | 4 | modarate |
| AGA | 89 | G>A | AAA | R89K | Neutral | Class C25 | TOLERATED | neutral | Reduced stability | BENIGN | Neutral | 1 | low |
| GCG | 91 | G>C | CCG | A91P | Neutral | Class C25 | AFFECT PROTEIN FUNCTION | neutral | Increased stability | BENIGN | Neutral | 1 | low |
| GCG | 91 | C>T | GTG | A91V | Neutral | Class C55 | AFFECT PROTEIN FUNCTION | neutral | Increased stability | BENIGN | Neutral | 2 | low |
| GCG | 92 | C>T | GTG | A92V | Neutral | Class C55 | TOLERATED | neutral | Reduced stability | BENIGN | Neutral | 2 | low |
| GGC | 94 | G>A | AGC | G94S | Neutral | Class C55 | AFFECT PROTEIN FUNCTION | neutral | Reduced stability | BENIGN | Neutral | 3 | modarate |
| TGT | 95 | T>A | AGT | C95S | Neutral | Class C65 | TOLERATED | neutral | Reduced stability | BENIGN | Neutral | 2 | low |
| CGG | 97 | G>A | CAG | R97Q | Neutral | Class C35 | TOLERATED | neutral | Reduced stability | POSSIBLY DAMAGING | Neutral | 2 | low |
| CAG | 100 | G>C, G>T | CAC, CAT | Q100H | Neutral | Class C15 | AFFECT PROTEIN FUNCTION | neutral | Reduced stability | BENIGN | Neutral | 2 | low |
| CGG | 104 | C>T | TGG | R104W | Deleterious | Class C65 | AFFECT PROTEIN FUNCTION | effect | Increased stability | PROBABLY DAMAGING | Neutral | 5 | modarate |
| CGG | 104 | G>A | CAG | R104Q | Neutral | Class C35 | AFFECT PROTEIN FUNCTION | neutral | Increased stability | PROBABLY DAMAGING | Neutral | 2 | low |
| GTG | 105 | T>G | GGG | V105G | Deleterious | Class C65 | AFFECT PROTEIN FUNCTION | effect | Reduced stability | PROBABLY DAMAGING | Neutral | 6 | high |
| GAC | 106 | G>A | AAC | D106N | Neutral | Class C15 | AFFECT PROTEIN FUNCTION | neutral | Reduced stability | PROBABLY DAMAGING | Neutral | 3 | modarate |
| CTC | 107 | C>T | TTC | L107F | Neutral | Class C15 | AFFECT PROTEIN FUNCTION | neutral | Increased stability | BENIGN | Neutral | 1 | low |
| TCA | 110 | T>A | ACA | S110T | Neutral | Class C55 | TOLERATED | neutral | Reduced stability | BENIGN | Neutral | 2 | low |
| CTC | 112 | C>A | ATC | L112I | Neutral | Class C0 | AFFECT PROTEIN FUNCTION | neutral | Increased stability | POSSIBLY DAMAGING | Neutral | 2 | low |
| CCA | 113 | C>T | TCA | P113S | Deleterious | Class C65 | AFFECT PROTEIN FUNCTION | neutral | Reduced stability | PROBABLY DAMAGING | Neutral | 5 | modarate |
| GAC | 115 | G>A | AAC | D115N | Neutral | Class C15 | AFFECT PROTEIN FUNCTION | neutral | Reduced stability | BENIGN | Neutral | 2 | low |
| CTG | 119 | T>G | CGG | L119R | Neutral | Class C65 | AFFECT PROTEIN FUNCTION | effect | Reduced stability | PROBABLY DAMAGING | Disease | 6 | high |
| AGG | 128 | G>A | AAG | R128K | Neutral | Class C25 | TOLERATED | neutral | Reduced stability | BENIGN | Neutral | 1 | low |
| AAG | 129 | G>T | AAT | K129N | Neutral | Class C65 | TOLERATED | neutral | Reduced stability | BENIGN | Neutral | 2 | low |
| GAC | 130 | G>A | AAC | D130N | Neutral | Class C15 | AFFECT PROTEIN FUNCTION | neutral | Reduced stability | BENIGN | Neutral | 2 | low |
| GAC | 130 | G>C | CAC | D130H | Neutral | Class C65 | AFFECT PROTEIN FUNCTION | neutral | Reduced stability | POSSIBLY DAMAGING | Neutral | 4 | modarate |
| CCA | 132 | C>A | ACA | P132T | Neutral | Class C35 | TOLERATED | neutral | Reduced stability | BENIGN | Neutral | 1 | low |
| CCA | 132 | C>A | CAA | P132Q | Neutral | Class C65 | TOLERATED | neutral | Reduced stability | BENIGN | Neutral | 2 | low |
| GCA | 138 | G>C | CCA | A138P | Neutral | Class C25 | TOLERATED | neutral | Reduced stability | POSSIBLY DAMAGING | Neutral | 2 | low |

Table 1.3: Mutation analysis result of RING0 region

| CODON | POSITION | CODON CHANGE | NEW CODON | MUTATION | proven | GV/GD | SIFT | SNAP | POLYPHEN2 | SDM | SNPs & GO | SCORE | INFERENCE |
| --- | --- | --- | --- | --- | --- | --- | --- | --- | --- | --- | --- | --- | --- |
| TCA | 141 | C>T | TTA | S141L | Neutral | Class C65 | AFFECT PROTEIN FUNCTION | effect | BENIGN | Increased stability | Neutral | 3 | modarate |
| AGC | 145 | G>A | AAC | S145N | Neutral | Class C45 | AFFECT PROTEIN FUNCTION | neutral | POSSIBLY DAMAGING | Reduced stability | Disease | 5 | modarate |
| GTG | 148 | G>T | TTG | V148L | Neutral | Class C25 | AFFECT PROTEIN FUNCTION | effect | POSSIBLY DAMAGING | Reduced stability | Disease | 5 | modarate |
| TGC | 150 | G>T | TTC | C150F | Deleterious | Class C65 | AFFECT PROTEIN FUNCTION | effect | PROBABLY DAMAGING | Reduced stability | Disease | 7 | high |
| GGC | 152 | G>A | GAC | G152D | Neutral | Class C65 | AFFECT PROTEIN FUNCTION | effect | BENIGN | Reduced stability | Disease | 5 | modarate |
| CCC | 153 | C>T | TCC | P153S | Deleterious | Class C65 | AFFECT PROTEIN FUNCTION | neutral | POSSIBLY DAMAGING | Reduced stability | Neutral | 5 | modarate |
| CCC | 153 | C>T | CTC | P153L | Deleterious | Class C65 | AFFECT PROTEIN FUNCTION | neutral | POSSIBLY DAMAGING | Reduced stability | Neutral | 5 | modarate |
| CCC | 153 | C>G | CGC | P153R | Deleterious | Class C65 | AFFECT PROTEIN FUNCTION | neutral | POSSIBLY DAMAGING | Reduced stability | Disease | 6 | high |
| AGA | 156 | G>T | ATA | R156I | Neutral | Class C65 | AFFECT PROTEIN FUNCTION | effect | POSSIBLY DAMAGING | Increased stability | Neutral | 4 | modarate |
| CAG | 158 | G>T | CAT | Q158H | Deleterious | Class C15 | AFFECT PROTEIN FUNCTION | neutral | PROBABLY DAMAGING | Reduced stability | Neutral | 4 | modarate |
| CCG | 159 | C>T | CTG | P159L | Deleterious | Class C65 | AFFECT PROTEIN FUNCTION | effect | PROBABLY DAMAGING | Increased stability | Neutral | 5 | modarate |
| GGA | 160 | G>A | GAA | G160E | Deleterious | Class C65 | AFFECT PROTEIN FUNCTION | effect | PROBABLY DAMAGING | Reduced stability | Disease | 7 | high |
| AAA | 161 | A>T | AAT | K161N | Deleterious | Class C65 | AFFECT PROTEIN FUNCTION | effect | PROBABLY DAMAGING | Reduced stability | Neutral | 6 | high |
| TGC | 166 | G>A | TAC | C166Y | Deleterious | Class C65 | AFFECT PROTEIN FUNCTION | effect | PROBABLY DAMAGING | Reduced stability | Disease | 7 | high |
| AGC | 167 | G>A | AAC | S167N | Neutral | Class C45 | AFFECT PROTEIN FUNCTION | neutral | BENIGN | Reduced stability | Neutral | 3 | modarate |
| CAA | 171 | A>T | CAT | Q171H | Deleterious | Class C15 | AFFECT PROTEIN FUNCTION | neutral | PROBABLY DAMAGING | Reduced stability | Neutral | 4 | modarate |
| GCA | 172 | G>T | TCA | A172S | Neutral | Class C65 | AFFECT PROTEIN FUNCTION | neutral | BENIGN | Reduced stability | Neutral | 3 | modarate |
| ACG | 173 | C>T | ATG | T173M | Deleterious | Class C65 | AFFECT PROTEIN FUNCTION | neutral | PROBABLY DAMAGING | Reduced stability | Neutral | 5 | modarate |
| TTG | 176 | G>C | TTC | L176F | Deleterious | Class C15 | AFFECT PROTEIN FUNCTION | effect | PROBABLY DAMAGING | Reduced stability | Neutral | 5 | modarate |
| GGT | 179 | G>T | TGT | G179C | Deleterious | Class C65 | AFFECT PROTEIN FUNCTION | neutral | PROBABLY DAMAGING | Reduced stability | Neutral | 5 | modarate |
| GAT | 185 | G>A | AAT | D185N | Deleterious | Class C15 | AFFECT PROTEIN FUNCTION | effect | PROBABLY DAMAGING | Reduced stability | Disease | 6 | high |
| GTT | 186 | G>A | ATT | V186I | Neutral | Class C25 | AFFECT PROTEIN FUNCTION | neutral | POSSIBLY DAMAGING | Reduced stability | Neutral | 3 | modarate |
| CCA | 189 | C>T | TCA | P189S | Deleterious | Class C65 | AFFECT PROTEIN FUNCTION | neutral | POSSIBLY DAMAGING | Reduced stability | Neutral | 5 | modarate |
| CGG | 191 | C>T | TGG | R191W | Deleterious | Class C65 | AFFECT PROTEIN FUNCTION | effect | PROBABLY DAMAGING | Reduced stability | Disease | 7 | high |
| CGG | 191 | G>A | CAG | R191Q | Deleterious | Class C35 | AFFECT PROTEIN FUNCTION | effect | PROBABLY DAMAGING | Reduced stability | Neutral | 5 | modarate |
| ATG | 192 | A>C | CTG | M192L | Neutral | Class C0 | AFFECT PROTEIN FUNCTION | effect | BENIGN | Reduced stability | Neutral | 3 | modarate |
| ATG | 192 | A>G | GTG | M192V | Neutral | Class C15 | AFFECT PROTEIN FUNCTION | neutral | BENIGN | Reduced stability | Neutral | 2 | low |
| GGT | 194 | GG>AT | ATT | G194I | Deleterious | Class C65 | AFFECT PROTEIN FUNCTION | effect | PROBABLY DAMAGING | Reduced stability | Disease | 7 | high |
| TGC | 196 | G>A | TAC | C196S | Deleterious | Class C65 | AFFECT PROTEIN FUNCTION | effect | PROBABLY DAMAGING | Reduced stability | Disease | 7 | high |
| TGC | 196 | G>C | TCC | C196Y | Deleterious | Class C65 | AFFECT PROTEIN FUNCTION | effect | PROBABLY DAMAGING | Reduced stability | Disease | 7 | high |
| CAC | 200 | C>G | CAG | H200Q | Neutral | Class C15 | TOLERATED | neutral | BENIGN | Reduced stability | Neutral | 1 | low |
| GGG | 203 | G>T | GTG | G203V | Deleterious | Class C65 | AFFECT PROTEIN FUNCTION | neutral | PROBABLY DAMAGING | Reduced stability | Neutral | 5 | modarate |
| GAA | 207 | G>A | AAA | E207K | Deleterious | Class C55 | AFFECT PROTEIN FUNCTION | effect | BENIGN | Reduced stability | Disease | 6 | high |
| TTT | 210 | T>G | TGT | F210C | Deleterious | Class C65 | AFFECT PROTEIN FUNCTION | effect | PROBABLY DAMAGING | Reduced stability | Disease | 7 | high |
| AAA | 211 | A>C | AAC | K211N | Deleterious | Class C65 | AFFECT PROTEIN FUNCTION | effect | PROBABLY DAMAGING | Reduced stability | Disease | 7 | high |
| TGT | 212 | G>A | TAT | C212Y | Deleterious | Class C65 | AFFECT PROTEIN FUNCTION | effect | PROBABLY DAMAGING | Reduced stability | Disease | 7 | high |
| TGT | 212 | T>G | GGT | C212G | Deleterious | Class C65 | AFFECT PROTEIN FUNCTION | effect | PROBABLY DAMAGING | Reduced stability | Disease | 7 | high |
| GGA | 213 | G>A | AGA | G213R | Deleterious | Class C65 | AFFECT PROTEIN FUNCTION | effect | POSSIBLY DAMAGING | Reduced stability | Disease | 7 | high |
| GGA | 213 | G>T | GTA | G213V | Deleterious | Class C65 | AFFECT PROTEIN FUNCTION | effect | POSSIBLY DAMAGING | Reduced stability | Neutral | 6 | high |
| GCA | 214 | C>A | GAA | A214E | Neutral | Class C65 | AFFECT PROTEIN FUNCTION | neutral | BENIGN | Reduced stability | Neutral | 3 | modarate |
| CAC | 215 | C>A | CAA | H215Q | Deleterious | Class C15 | AFFECT PROTEIN FUNCTION | effect | PROBABLY DAMAGING | Reduced stability | Disease | 6 | high |
| CCC | 216 | C>T | TCC | P216S | Deleterious | Class C65 | AFFECT PROTEIN FUNCTION | neutral | BENIGN | Reduced stability | Neutral | 4 | modarate |

Table 1.4: Mutation analysis results of RING1 region

| CODON | POSITION | CODON CHANGE | NEW CODON | MUTATION | PROVEN | GV/GD | SIFT | SNAP | SDM | POLYPHEN2 | SNPs & GO | score | Inference |
| --- | --- | --- | --- | --- | --- | --- | --- | --- | --- | --- | --- | --- | --- |
| GAA | 221 | G>A | AAA | E221K | Deleterious | Class C55 | AFFECT PROTEIN FUNCTION | effect | Reduced stability | BENIGN | Disease | 6 | HIGH |
| ACA | 222 | A>G | GAC | T222A | Neutral | Class C55 | TOLERATED | neutral | Reduced stability | BENIGN | Neutral | 2 | LOW |
| TTG | 226 | G>T | TTT | L226F | Deleterious | Class C15 | AFFECT PROTEIN FUNCTION | effect | Reduced stability | POSSIBLY DAMAGING | Disease | 6 | HIGH |
| CTG | 228 | C>A | ATG | L228M | Neutral | Class C0 | AFFECT PROTEIN FUNCTION | neutral | Reduced stability | PROBABLY DAMAGING | Disease | 4 | Modarate |
| ATC | 229 | A>T | TTC | I229F | Deleterious | Class C15 | AFFECT PROTEIN FUNCTION | effect | Reduced stability | BENIGN | Disease | 5 | Modarate |
| GCA | 230 | G>A | ACA | A230T | Neutral | Class C55 | TOLERATED | neutral | Reduced stability | BENIGN | Neutral | 2 | LOW |
| CGG | 234 | G>A | CAG | R234Q | Neutral | Class C35 | TOLERATED | neutral | Reduced stability | POSSIBLY DAMAGING | Neutral | 2 | LOW |
| ACT | 237 | A>G | GCT | T237A | Neutral | Class C55 | TOLERATED | effect | Reduced stability | BENIGN | Neutral | 3 | Modarate |
| TGC | 238 | C>G | TGG | C238W | Deleterious | Class C65 | AFFECT PROTEIN FUNCTION | effect | Reduced stability | PROBABLY DAMAGING | Disease | 7 | HIGH |
| ACG | 240 | C>T | ATG | T240M | Neutral | Class C65 | TOLERATED | effect | Reduced stability | POSSIBLY DAMAGING | Neutral | 4 | Modarate |
| ACG | 240 | C>G | AGG | T240R | Neutral | Class C65 | AFFECT PROTEIN FUNCTION | effect | Reduced stability | POSSIBLY DAMAGING | Disease | 6 | HIGH |
| GAC | 243 | G>A | AAC | D243N | Deleterious | Class C15 | AFFECT PROTEIN FUNCTION | effect | Reduced stability | POSSIBLY DAMAGING | Disease | 6 | HIGH |
| GTC | 244 | G>A | ATC | V244I | Neutral | Class C25 | TOLERATED | neutral | Increased stability | BENIGN | Neutral | 0 | LOW |
| CAG | 252 | A>G | CGG | Q252R | Neutral | Class C35 | TOLERATED | effect | Reduced stability | POSSIBLY DAMAGING | Neutral | 3 | Modarate |
| TGC | 253 | G>A | TAC | C253Y | Deleterious | Class C65 | AFFECT PROTEIN FUNCTION | effect | Reduced stability | PROBABLY DAMAGING | Disease | 7 | HIGH |
| TCC | 255 | C>T | TTC | S255F | Deleterious | Class C65 | TOLERATED | neutral | Reduced stability | BENIGN | Neutral | 3 | Modarate |
| CGC | 256 | G>T | CTC | R256L | Deleterious | Class C65 | TOLERATED | effect | Reduced stability | PROBABLY DAMAGING | Disease | 6 | HIGH |
| CGC | 256 | C>T | TGC | R256C | Deleterious | Class C65 | AFFECT PROTEIN FUNCTION | effect | Reduced stability | PROBABLY DAMAGING | Disease | 7 | HIGH |
| GTG | 258 | G>A | ATG | V258M | Deleterious | Class C15 | TOLERATED | neutral | Reduced stability | PROBABLY DAMAGING | Neutral | 3 | Modarate |
| TGC | 260 | G>T | TTC | C260F | Deleterious | Class C65 | AFFECT PROTEIN FUNCTION | effect | Reduced stability | PROBABLY DAMAGING | Disease | 7 | HIGH |
| GAC | 262 | G>C | CAC | D262H | Deleterious | Class C65 | AFFECT PROTEIN FUNCTION | neutral | Reduced stability | POSSIBLY DAMAGING | Disease | 6 | HIGH |
| TTA | 266 | T>G | GTA | L266V | Neutral | Class C25 | TOLERATED | neutral | Reduced stability | POSSIBLY DAMAGING | Neutral | 2 | LOW |
| ACA | 270 | A>G | GCA | T270A | Neutral | Class C55 | AFFECT PROTEIN FUNCTION | neutral | Increased stability | PROBABLY DAMAGING | Neutral | 3 | Modarate |
| AGA | 271 | G>A | AAA | R271K | Neutral | Class C25 | TOLERATED | effect | Reduced stability | BENIGN | Neutral | 2 | LOW |
| AGA | 271 | A>T | AGT | R271S | Deleterious | Class C65 | TOLERATED | effect | Reduced stability | PROBABLY DAMAGING | Disease | 6 | HIGH |
| CTC | 272 | C>A | ATC | L272I | Neutral | Class C0 | TOLERATED | effect | Reduced stability | PROBABLY DAMAGING | Neutral | 3 | Modarate |
| AAT | 273 | A>G | AGT | N273S | Deleterious | Class C45 | TOLERATED | neutral | Reduced stability | PROBABLY DAMAGING | Neutral | 4 | Modarate |
| GAT | 274 | G>A | AAT | D274N | Deleterious | Class C15 | AFFECT PROTEIN FUNCTION | effect | Reduced stability | BENIGN | Neutral | 4 | Modarate |
| GAT | 274 | G>C | CAT | D274H | Deleterious | Class C65 | AFFECT PROTEIN FUNCTION | effect | Reduced stability | PROBABLY DAMAGING | Disease | 7 | HIGH |
| CGG | 275 | C>T | TGG | R275W | Deleterious | Class C65 | AFFECT PROTEIN FUNCTION | effect | Reduced stability | PROBABLY DAMAGING | Disease | 7 | HIGH |
| CGG | 275 | G>C | CCG | R275P | Deleterious | Class C65 | TOLERATED | effect | Reduced stability | PROBABLY DAMAGING | Disease | 6 | HIGH |
| CGG | 275 | G>T | CTG | R275L | Deleterious | Class C65 | TOLERATED | effect | Reduced stability | PROBABLY DAMAGING | Disease | 6 | HIGH |
| TTT | 277 | T>G | TGT | F277C | Deleterious | Class C65 | AFFECT PROTEIN FUNCTION | effect | Reduced stability | PROBABLY DAMAGING | Disease | 7 | HIGH |
| GAC | 280 | G>A | AAC | D280N | Neutral | Class C15 | TOLERATED | effect | Reduced stability | POSSIBLY DAMAGING | Neutral | 3 | Modarate |
| CCT | 281 | C>T | TCT | P281S | Deleterious | Class C65 | TOLERATED | neutral | Reduced stability | POSSIBLY DAMAGING | Neutral | 4 | Modarate |
| CAA | 282 | A>C | CAC | Q282H | Neutral | Class C15 | TOLERATED | neutral | Reduced stability | BENIGN | Neutral | 1 | LOW |
| CTT | 283 | T>G | CGT | L283R | Deleterious | Class C65 | AFFECT PROTEIN FUNCTION | effect | Reduced stability | PROBABLY DAMAGING | Disease | 7 | HIGH |
| CTT | 283 | T>C | CCT | L283P | Deleterious | Class C65 | AFFECT PROTEIN FUNCTION | effect | Reduced stability | PROBABLY DAMAGING | Disease | 7 | HIGH |
| GGC | 284 | G>C | CGC | G284R | Deleterious | Class C65 | AFFECT PROTEIN FUNCTION | effect | Reduced stability | PROBABLY DAMAGING | Disease | 7 | HIGH |
| TCC | 286 | C>G | TGC | S286C | Deleterious | Class C65 | TOLERATED | effect | Increased stability | PROBABLY DAMAGING | Neutral | 4 | Modarate |
| TGT | 289 | T>G | GGT | C289G | Deleterious | Class C65 | AFFECT PROTEIN FUNCTION | effect | Reduced stability | PROBABLY DAMAGING | Disease | 7 | HIGH |
| GTG | 290 | T>A | GAG | V290E | Neutral | Class C65 | TOLERATED | effect | Reduced stability | PROBABLY DAMAGING | Neutral | 4 | Modarate |
| GCT | 291 | G>A | ACT | A291T | Deleterious | Class C55 | AFFECT PROTEIN FUNCTION | effect | Reduced stability | PROBABLY DAMAGING | Neutral | 6 | HIGH |
| GCT | 291 | G>T | TCT | A291S | Neutral | Class C65 | TOLERATED | effect | Reduced stability | POSSIBLY DAMAGING | Neutral | 4 | Modarate |
| GGC | 292 | G>A | AGC | G292S | Deleterious | Class C55 | TOLERATED | effect | Reduced stability | BENIGN | Neutral | 4 | Modarate |
| GGC | 292 | G>A | GAC | G292D | Deleterious | Class C65 | TOLERATED | effect | Reduced stability | PROBABLY DAMAGING | Disease | 6 | HIGH |
| CCC | 294 | C>T | CTC | P294L | Deleterious | Class C65 | TOLERATED | neutral | Reduced stability | PROBABLY DAMAGING | Neutral | 4 | Modarate |
| AAC | 295 | C>A | AAA | N295K | Deleterious | Class C65 | TOLERATED | effect | Reduced stability | POSSIBLY DAMAGING | Neutral | 5 | Modarate |
| ATT | 298 | A>T | TTT | I298F | Deleterious | Class C15 | TOLERATED | effect | Reduced stability | PROBABLY DAMAGING | Disease | 5 | Modarate |
| ATT | 298 | A>C | CTT | I298L | Neutral | Class C0 | TOLERATED | effect | Increased stability | POSSIBLY DAMAGING | Disease | 4 | Modarate |
| ATT | 298 | T>G | AGT | I298S | Deleterious | Class C65 | AFFECT PROTEIN FUNCTION | effect | Reduced stability | PROBABLY DAMAGING | Disease | 7 | HIGH |
| GAG | 300 | A>G | GGG | E300G | Deleterious | Class C65 | AFFECT PROTEIN FUNCTION | effect | Reduced stability | PROBABLY DAMAGING | Disease | 7 | HIGH |
| CTC | 301 | C>T | TTC | L301F | Neutral | Class C15 | TOLERATED | effect | Reduced stability | PROBABLY DAMAGING | Neutral | 3 | Modarate |
| CAC | 303 | C>T | TAC | H303Y | Deleterious | Class C65 | TOLERATED | effect | Reduced stability | PROBABLY DAMAGING | Disease | 6 | HIGH |
| GGA | 308 | G>A | GAA | G308E | Deleterious | Class C65 | TOLERATED | neutral | Reduced stability | PROBABLY DAMAGING | Neutral | 4 | Modarate |
| GAG | 310 | G>C | GAC | E310D | Neutral | Class C35 | TOLERATED | neutral | Reduced stability | BENIGN | Neutral | 1 | LOW |
| CGG | 311 | G>T | TGG | Q311H | Deleterious | Class C15 | TOLERATED | effect | Reduced stability | PROBABLY DAMAGING | Disease | 5 | Modarate |
| CGG | 314 | C>T | CGG | R314W | Deleterious | Class C65 | AFFECT PROTEIN FUNCTION | effect | Reduced stability | PROBABLY DAMAGING | Disease | 7 | HIGH |
| CAG | 317 | A>G | CGG | Q317R | Neutral | Class C35 | TOLERATED | neutral | Increased stability | BENIGN | Neutral | 0 | LOW |
| GCA | 320 | G>T | TCA | A320S | Neutral | Class C65 | TOLERATED | neutral | Reduced stability | POSSIBLY DAMAGING | Neutral | 3 | Modarate |
| GTC | 324 | G>T | TTC | V324F | Deleterious | Class C45 | TOLERATED | effect | Increased stability | PROBABLY DAMAGING | Disease | 5 | Modarate |
| CTG | 325 | C>A | ATG | L325M | Neutral | Class C0 | TOLERATED | effect | Increased stability | PROBABLY DAMAGING | Neutral | 2 | LOW |
| CAG | 326 | G>T | CAT | Q326H | Deleterious | Class C15 | AFFECT PROTEIN FUNCTION | neutral | Reduced stability | BENIGN | Neutral | 3 | Modarate |
| ATG | 327 | G>A | ATA | M327I | Neutral | Class C0 | TOLERATED | neutral | Reduced stability | BENIGN | Neutral | 1 | LOW |
| GGG | 328 | G>T | GTG | G328V | Deleterious | Class C65 | TOLERATED | effect | Reduced stability | PROBABLY DAMAGING | Disease | 6 | HIGH |
| GGG | 328 | G>A | GAG | G328E | Deleterious | Class C65 | TOLERATED | effect | Reduced stability | PROBABLY DAMAGING | Disease | 6 | HIGH |

Table 1.5: Mutation analysis result of REP region

| CODON | POSITION | AMINO ACID CHANGE | NEW CODON | Mutation | Domain | SNPs&GO | SNAP2 | Polyphen2 | SIFT | SDM | Proven | Aline GV&GD | score |  |
| --- | --- | --- | --- | --- | --- | --- | --- | --- | --- | --- | --- | --- | --- | --- |
| GCC | 379 | C>T | GTC | A379V | REP | Neutral | neutral | BENIGN | TOLERATED | Reduced stability | Neutral | Class C55 | 2 | Low |
| GTA | 380 | G>C | CTA | V380I | REP | Neutral | neutral | BENIGN | TOLERATED | Increased stability | Neutral | Class C25 | 0 | Low |
| GGA | 385 | G>A | GAA | G385E | REP | Neutral | effect | BENIGN | TOLERATED | Reduced stability | Neutral | Class C65 | 3 | Modarate |
| ACT | 387 | C>G | AGT | T387S | REP | Neutral | neutral | BENIGN | TOLERATED | Reduced stability | Neutral | Class C55 | 2 | Low |
| CAG | 389 | G>T | CAT | Q389H | REP | Neutral | neutral | POSSIBLY DAMAGING | AFFECT PROTEIN FUNCTION | Reduced stability | Neutral | Class C15 | 3 | Modarate |
| AGA | 392 | A>G | GGA | R392G | REP | Neutral | effect | BENIGN | AFFECT PROTEIN FUNCTION | Reduced stability | Neutral | Class C65 | 4 | Modarate |
| GTC | 393 | G>A | ATC | V393I | REP | Neutral | neutral | POSSIBLY DAMAGING | TOLERATED | Reduced stability | Neutral | Class C25 | 2 | Low |
| GAT | 394 | A>G | GGT | D394G | REP | Neutral | effect | BENIGN | TOLERATED | Reduced stability | Deleterious | Class C65 | 4 | modarate |
| GAT | 394 | G>A | AAT | D394N | REP | Neutral | effect | POSSIBLY DAMAGING | TOLERATED | Reduced stability | Deleterious | Class C15 | 4 | modarate |
| GAA | 395 | A>C | GCA | E395A | REP | Neutral | neutral | BENIGN | TOLERATED | Increased stability | Neutral | Class C65 | 1 | Low |
| AGA | 396 | A>G | GGA | R396G | REP | Neutral | neutral | BENIGN | TOLERATED | Reduced stability | Neutral | Class C65 | 2 | Low |
| GCC | 397 | C>T | ACC | A397V | REP | Neutral | neutral | PROBABLY DAMAGING | TOLERATED | Reduced stability | Neutral | Class C55 | 3 | modarate |
| GCC | 398 | G>A | ACC | A398T | REP | Neutral | neutral | PROBABLY DAMAGING | TOLERATED | Reduced stability | Neutral | Class C55 | 3 | modarate |
| CGT | 402 | C>T | TGT | R402G | REP | Disease | effect | PROBABLY DAMAGING | TOLERATED | Reduced stability | Deleterious | Class C65 | 6 | High |
| GAA | 404 | G>A | AAA | E404K | REP | Neutral | effect | PROBABLY DAMAGING | AFFECT PROTEIN FUNCTION | Reduced stability | Deleterious | Class C55 | 6 | High |
| TCC | 407 | T>C | CCC | S407P | REP | Disease | effect | PROBABLY DAMAGING | AFFECT PROTEIN FUNCTION | Reduced stability | Deleterious | Class C65 | 7 | High |
| TCC | 407 | C>G | TGC | S407C | REP | Neutral | effect | PROBABLY DAMAGING | AFFECT PROTEIN FUNCTION | Increased stability | Deleterious | Class C66 | 5 | modarate |
| TCC | 407 | C>T | TTC | S407F | REP | Disease | effect | PROBABLY DAMAGING | AFFECT PROTEIN FUNCTION | Reduced stability | Deleterious | Class C67 | 7 | High |
| AAA | 408 | A>C | AAC | K408N | REP | Neutral | neutral | PROBABLY DAMAGING | TOLERATED | Reduced stability | Neutral | Class C68 | 3 | modarate |
| GAA | 409 | G>A | AAA | E409K | REP | Neutral | neutral | PROBABLY DAMAGING | TOLERATED | Reduced stability | Neutral | Class C55 | 3 | modarate |

Table 1.6: Mutation analysis results of IBR region

| Mutation | Domain | CODON | POSITION | CODON CHANGE | NEW CODON | MUTATION | AA change | SNAP (Predicted Effect) | Aline GV-GD (Prediction) | Proven (Prediction (cutoff= -2.5)) | SNPS & GO (Effect) | SDM | SIFT | Polyphen2 (Effect) | Prediction |  |
| --- | --- | --- | --- | --- | --- | --- | --- | --- | --- | --- | --- | --- | --- | --- | --- | --- |
| G329S | IBR | GGC | 329 | G>A | AGC | G329S | G>S | effect | Class C55 | Deleterious | Neutral | Reduced stability | TOLERATED | Probably damaging | 5 | MODARATE |
| G329D | IBR | GGC | 329 | G>A | GAC | G329D | G>D | effect | Class C65 | Deleterious | Neutral | Reduced stability | TOLERATED | Probably damaging | 5 | MODARATE |
| V330M | IBR | GTG | 330 | G>A | ATG | V330M | V>M | effect | Class C15 | Neutral | Neutral | Reduced stability | AFFECT PROTEIN FUNCTION | Probably damaging | 4 | MODARATE |
| R334H | IBR | CGC | 334 | G>A | CAC | R334H | R>H | effect | Class C25 | Neutral | Neutral | Reduced stability | TOLERATED | Benign | 2 | LOW |
| R334C | IBR | CGC | 334 | C>T | TGC | R334C | R>C | effect | Class C65 | Deleterious | Neutral | Reduced stability | AFFECT PROTEIN FUNCTION | Probably damaging | 6 | HIGH |
| P335S | IBR | CCT | 335 | C>T | TCT | P335S | P>S | effect | Class C65 | Deleterious | Disease | Reduced stability | TOLERATED | Probably damaging | 6 | HIGH |
| P335L | IBR | CCT | 335 | C>T | CTT | P335L | P>L | effect | Class C65 | Deleterious | Disease | Reduced stability | AFFECT PROTEIN FUNCTION | Probably damaging | 7 | HIGH |
| G338V | IBR | GGA | 338 | G>T | GTA | G337V | G>V | effect | Class C65 | Deleterious | Disease | Reduced stability | TOLERATED | Probably damaging | 6 | HIGH |
| A339S | IBR | GCG | 339 | G>T | TCG | A338S | A>S | effect | Class C65 | Neutral | Neutral | Reduced stability | TOLERATED | Probably damaging | 4 | MODARATE |
| G340W | IBR | GGG | 340 | G>T | TGG | G339W | G>W | effect | Class C65 | Deleterious | Disease | Reduced stability | AFFECT PROTEIN FUNCTION | Probably damaging | 7 | HIGH |
| P343Q | IBR | CCG | 343 | C>A | CAG | P343Q | P>Q | effect | Class C65 | Deleterious | Neutral | Reduced stability | TOLERATED | Probably damaging | 5 | MODARATE |
| P343L | IBR | CCG | 343 | C>T | CTG | P343L | P>L | neutral | Class C65 | Deleterious | Neutral | Reduced stability | TOLERATED | Probably damaging | 4 | MODARATE |
| E344K | IBR | GAG | 344 | G>A | AAG | E344K | E>K | effect | Class C55 | Neutral | Neutral | Reduced stability | TOLERATED | Possibly damaging | 4 | MODARATE |
| D346N | IBR | GAC | 346 | G>A | AAC | D346N | D>N | neutral | Class C15 | Neutral | Neutral | Reduced stability | TOLERATED | Possibly damaging | 2 | LOW |
| Q347H | IBR | CAG | 347 | G>C, G>T | CAC,CAT | Q347H | Q>H | neutral | Class C15 | Neutral | Neutral | Reduced stability | AFFECT PROTEIN FUNCTION | Possibly damaging | 3 | MODARATE |
| T351P | IBR | ACC | 351 | A>C | CCC | T351P | T>P | effect | Class C35 | Neutral | Neutral | Reduced stability | TOLERATED | Probably damaging | 3 | MODARATE |
| E353K | IBR | GAA | 353 | G>A | AAA | E353K | E>K | effect | Class C55 | Neutral | Neutral | Reduced stability | TOLERATED | Probably damaging | 4 | MODARATE |
| G354R | IBR | GGG | 354 | G>C | CGG | G354R | G>R | neutral | Class C65 | Neutral | Neutral | Reduced stability | TOLERATED | Benign | 2 | LOW |
| G359D | IBR | GGC | 359 | G>A | GAC | G359D | G>D | effect | Class C65 | Deleterious | Disease | Reduced stability | TOLERATED | Probably damaging | 6 | HIGH |
| C360F | IBR | TGT | 360 | G>T | TTT | C360F | C>F | effect | Class C65 | Deleterious | Disease | Reduced stability | AFFECT PROTEIN FUNCTION | Probably damaging | 7 | HIGH |
| F364V | IBR | TTC | 364 | T>G | GTC | F364V | F>V | effect | Class C45 | Deleterious | Disease | Reduced stability | AFFECT PROTEIN FUNCTION | Probably damaging | 7 | HIGH |
| F364L | IBR | TTC | 364 | C>A | TTA | F364L | F>L | effect | Class C15 | Deleterious | Disease | Reduced stability | AFFECT PROTEIN FUNCTION | Probably damaging | 6 | HIGH |
| R366Q | IBR | CGG | 366 | G>A | CAG | R366Q | R>Q | effect | Class C35 | Deleterious | Disease | Reduced stability | TOLERATED | Probably damaging | 5 | MODARATE |
| R366W | IBR | GAA | 370 | G>A | AAA | R366W | R>W | effect | Class C65 | Deleterious | Disease | Reduced stability | TOLERATED | Probably damaging | 6 | HIGH |
| E370K | IBR | ACG | 371 | G>A | ACA | E371K | E>K | effect | Class C55 | Neutral | Neutral | Reduced stability | TOLERATED | Possibly damaging | 4 | MODARATE |
| H373D | IBR | CAT | 373 | C>G | GAT | H373D | H>D | effect | Class C65 | Deleterious | Disease | Reduced stability | AFFECT PROTEIN FUNCTION | Benign | 6 | HIGH |
| H373R | IBR | CAT | 373 | A>G | CGT | H373R | H>R | effect | Class C25 | Deleterious | Disease | Reduced stability | TOLERATED | Probably damaging | 5 | MODARATE |
| S378G | IBR | AGT | 378 | A>G | GGT | S378G | S>G | neutral | Class C55 | Neutral | Neutral | Reduced stability | TOLERATED | Benign | 2 | LOW |

Table 1.7: Mutation analysis results of RING2 region

| CODON | POSITION | CODON CHANGE | NEW CODON | MUTATION | PROVEN | GV/GD | SIFT | SNAP | SDM | POLYPHEN2 | SNPS & Go | score |  |
| --- | --- | --- | --- | --- | --- | --- | --- | --- | --- | --- | --- | --- | --- |
| ACC | 415 | C>A | AAC | T415N | Deleterious | Class C55 | AFFECT PROTEIN FUNCTION | effect | Reduced stability | PROBABLY DAMAGING | Disease | 7 | high |
| CCC | 417 | C>T | CTC | P417L | Deleterious | Class C65 | TOLERATED | effect | Increased stability | PROBABLY DAMAGING | Neutral | 5 | modarate |
| TGT | 418 | T>C | CGT | C418R | Deleterious | Class C65 | AFFECT PROTEIN FUNCTION | effect | Reduced stability | PROBABLY DAMAGING | Disease | 7 | high |
| CCC | 419 | C>T | CTC | P419L | Deleterious | Class C65 | AFFECT PROTEIN FUNCTION | effect | Reduced stability | PROBABLY DAMAGING | Disease | 7 | high |
| CGC | 420 | C>T | TGC | R420C | Deleterious | Class C65 | TOLERATED | effect | Reduced stability | PROBABLY DAMAGING | Neutral | 5 | modarate |
| CGC | 420 | G>A | CAC | R420H | Neutral | Class C25 | AFFECT PROTEIN FUNCTION | effect | Reduced stability | PROBABLY DAMAGING | Neutral | 4 | modarate |
| GAA | 426 | A>G | GAG | E426G | Deleterious | Class C65 | AFFECT PROTEIN FUNCTION | effect | Reduced stability | PROBABLY DAMAGING | Disease | 7 | high |
| GGA | 429 | G>A | GAA | G429E | Deleterious | Class C65 | TOLERATED | effect | Reduced stability | PROBABLY DAMAGING | Disease | 6 | high |
| GGC | 430 | G>A | GAC | G430D | Deleterious | Class C65 | AFFECT PROTEIN FUNCTION | effect | Reduced stability | PROBABLY DAMAGING | Disease | 7 | high |
| TGC | 431 | G>C | TCC | C431F | Deleterious | Class C65 | AFFECT PROTEIN FUNCTION | effect | Reduced stability | PROBABLY DAMAGING | Disease | 7 | high |
| ATG | 432 | G>A | ATA | M432I | Deleterious | Class C0 | TOLERATED | effect | Reduced stability | PROBABLY DAMAGING | Neutral | 4 | modarate |
| CCG | 437 | C>T | CTG | P437L | Neutral | Class C65 | AFFECT PROTEIN FUNCTION | effect | Reduced stability | PROBABLY DAMAGING | Neutral | 5 | modarate |
| CAG | 438 | C>A | AAG | Q438K | Neutral | Class C45 | TOLERATED | effect | Reduced stability | BENIGN | Neutral | 3 | modarate |
| TGC | 441 | T>C | CGC | C441R | Deleterious | Class C65 | AFFECT PROTEIN FUNCTION | effect | Reduced stability | PROBABLY DAMAGING | Disease | 7 | high |
| GAG | 444 | G>A | AAG | E444K | Neutral | Class C55 | AFFECT PROTEIN FUNCTION | effect | Reduced stability | PROBABLY DAMAGING | Disease | 6 | high |
| GAG | 444 | G>C | CAG | E444Q | Neutral | Class C25 | AFFECT PROTEIN FUNCTION | effect | Reduced stability | PROBABLY DAMAGING | Neutral | 4 | modarate |
| TGT | 449 | T>C | CGT | C449R | Deleterious | Class C65 | AFFECT PROTEIN FUNCTION | effect | Reduced stability | PROBABLY DAMAGING | Disease | 7 | high |
| TGT | 449 | G>T | TTT | C449F | Deleterious | Class C65 | AFFECT PROTEIN FUNCTION | effect | Reduced stability | PROBABLY DAMAGING | Disease | 7 | high |
| TGG | 453 | G>T | TTG | W453L | Deleterious | Class C55 | TOLERATED | effect | Reduced stability | PROBABLY DAMAGING | Disease | 6 | high |
| CGC | 455 | C>A | AGC | R455S | Deleterious | Class C65 | AFFECT PROTEIN FUNCTION | effect | Reduced stability | PROBABLY DAMAGING | Disease | 7 | high |
| GTC | 456 | G>A | ATC | V456I | Neutral | Class C25 | TOLERATED | neutral | Increased stability | BENIGN | Neutral | 0 | low |
| ATG | 458 | A>C | CTG | M458L | Neutral | Class C0 | TOLERATED | neutral | Increased stability | BENIGN | Neutral | 0 | low |
| CAC | 461 | A>G | CGC | H461R | Deleterious | Class C25 | AFFECT PROTEIN FUNCTION | effect | Reduced stability | PROBABLY DAMAGING | Disease | 6 | high |
| TGG | 462 | G>T | TTG | W462L | Deleterious | Class C55 | AFFECT PROTEIN FUNCTION | effect | Reduced stability | PROBABLY DAMAGING | Disease | 7 | high |
| GAC | 464 | G>A | AAC | D464N | Neutral | Class C15 | TOLERATED | effect | Reduced stability | POSSIBLY DAMAGING | Neutral | 3 | modarate |
| GAC | 464 | A>T | GTC | D464V | Deleterious | Class C65 | AFFECT PROTEIN FUNCTION | effect | Reduced stability | PROBABLY DAMAGING | Neutral | 6 | high |
